# Mapping the microbial diversity and natural resistome of North Antarctica soils

**DOI:** 10.1101/2021.05.05.442734

**Authors:** Andrés E. Marcoleta, Macarena A Varas, José Costa, Johanna Rojas-Salgado, Patricio Arros, Camilo Berríos-Pastén, Sofía Tapia, Daniel Silva, José Fierro, Nicolás Canales, Francisco P Chávez, Alexis Gaete, Mauricio González, Miguel A. Allende, Rosalba Lagos

## Abstract

The rising of multiresistant bacterial pathogens is currently one of the most critical threats to global health, demanding a better understanding of the origin and spread of antibiotic resistance. In this regard, the resistome hosted by the microbiota from natural and remote environments remains poorly explored. Moreover, little is known about the availability of antimicrobial resistance genes (ARGs) from these environments to be disseminated through horizontal transfer, potentially mediating the rise of novel resistance factors among clinically relevant pathogens. In this context, the North Antarctica soils are attractive ecosystems to study due to the presence of a microbiota naturally adapted to thrive in harsh conditions, including potential factors to resist natural toxic substances. In this work, we evaluated the antibiotic resistance of bacteria isolated from soils collected in humanized and non-intervened areas of North Antarctica. We identified resistance to a wide array of antibiotics, with isolates harboring up to 10 simultaneous resistances, mainly native *Pseudomonas*. Genomic analysis revealed the presence of a wide array of genes encoding efflux pumps but the lack of genes explaining some of the resistance phenotypes, suggesting novel uncharacterized mechanisms. Also, using 16S rRNA amplicon and shotgun metagenome sequencing, we explored the microbial diversity in the sampled soils and evaluated the presence of ARGs and their host microbiota. High microbial diversity was found in all the sites, with Proteobacteria, Bacteroidota, Acidobacteriota, and Verrucomicrobiota being the most abundant Phyla, while *Candidatus Udaeobacter*, RB41, *Polaromonas*, and *Ferruginibacter* the most abundant genera. We identified hundreds of genes potentially conferring resistance to more than 15 drug classes, both by short reads analyses and ARG detection among assembled contigs and MAGs obtained combining short and long-read sequence data. *Polaromonas, Pseudomonas, Streptomyces, Variovorax, Bhurkolderia*, and *Gemmatimonas* were the main host taxa of the identified ARGs. Part of these ARGs was found inside predicted plasmids, including a putative OXA-like beta-lactamase from *Polaromonas* harboring the key conserved residues of this kind of enzyme and a conserved predicted protein structure. All this evidence indicates that microbial communities from North Antarctica soil have a highly diverse natural resistome, part of it located inside mobile genetic elements, which would act as a source of novel ARGs.

## INTRODUCTION

Currently, antimicrobial therapies’ efficacy is rapidly declining worldwide, mainly because of the increasing detection of multiresistant bacterial strains. Thus, antimicrobial resistance is now among the most pressing health crises of the 21^st^ Century, demanding urgent actions to prevent the dissemination of resistant pathogens and encouraging the research to understand the origin, evolution, and spread of antibiotic resistance (Hutchings et al., 2019; Smith et al., 2015). As for other global health issues, this crisis should be faced following the so-called “One-health” framework, which stresses that human health is interconnected with animal health and the environment. In this regard, although antimicrobial resistance has been studied thoroughly in humans, livestock, and to a lesser extent in contaminated environments, considerably less is known regarding the resistome (i.e. pool of resistance factors) present in natural and remote environments.

Recent reports have evidenced the presence, in remote areas, of antibiotic resistance genes typically detected among bacterial pathogens infecting humans. This last is the case of the bla_NDM-1_ carbapenemase, commonly associated with multiresistant *Enterobacteriaceae* causing outbreaks, which gene was detected by quantitative PCR among soil samples collected from the High Arctic (northern Norway) (McCann et al., 2019). In this study, a gradient of potential human and wildlife impacts across zones correlated positively with the presence of this and other foreign resistance genes. A similar observation was made by Tan et al., studying Arctic marine sediments collected from the Bering Sea, which found significant correlations between the copy number of resistance genes and the human mitochondrial gene Hmt as a marker of human contamination, as well as with the concentration of different antibiotics found in the sediments. Additionally, previous reports suggest that human activity and animal carriage would have promoted the presence of antibiotic-resistant bacteria isolated from freshwater in King George Island, Antarctica (Jara et al., 2020). This evidence suggests a concerning spread of resistance genes from pathogens contaminating diverse natural and remote environments, possibly through carriage by migrating birds and animals connecting these environments with humanized areas. Also, it points out the potential influence of anthropogenic activities on the presence of antibiotic-resistant bacteria in the environment.

On the other hand, it was also documented the presence of potential resistance genes from autochthonous bacteria living in the wild, even in remote non-intervened areas, including soils and marine sediments from the Arctic, and soils within the undisturbed Mackay Glacier region in Antarctica (Goethem et al., 2018; McCann et al., 2019; Tan et al., 2018). This line of evidence suggests that the development of antibiotic resistance is a natural phenomenon occurring widespread on our planet that, as for other metabolic features, developed through billions of years of evolution (Blair et al., 2015; Forsberg et al., 2012; Hutchings et al., 2019). Furthermore, most clinically used antibiotics derive from antibiotic-producing microorganisms that naturally expose other species to these substances in their local environment. This would favor the development and selection of resistance in both the affected environmental species and the producer organisms. Also, proteins such as efflux pumps, allowing environmental bacteria to deal with toxic compounds produced on their niche, could be leveraged by pathogens to resist clinically used antibiotics sharing structural or chemical similarity with these natural compounds. Thus, a wide diversity of genes from microbes living in natural environments, conferring resistance to potentially all the known antibiotics, is yet to be characterized. However, there is limited information regarding the mobility of these genes and possible dissemination mechanisms to reach pathogenic bacteria. In particular, less is known regarding the natural resistome found in extreme environments, where the resident microbiota would have developed remarkable adaptations to thrive in harsh conditions, such as the presence of antimicrobials and other toxic substances.

In this work, we studied the bacterial communities living in soils from different locations in North Antarctica as model ecosystems to investigate the presence of antibiotic-resistant bacteria and resistance genes, comparing areas intervened by human activities with protected areas of restricted access without evident human intervention. We tested the resistance to a variety of antibiotics of over 200 psychrotolerant Antarctic isolates, identifying a number of multiresistant strains. Also, we performed genome analysis of two of them, searching for the genetic basis of the resistance phenotype, which suggested the presence of new resistance mechanisms. Furthermore, we conducted metagenomic approaches combining illumina and nanopore sequencing to study the microbial diversity and the presence of resistance genes in humanized and non-intervened zones from North Antarctica, determining which bacteria host these genes, and searching for resistance factors encoded in plasmids.

## METHODS

### Antarctic soil sample collection and bacterial isolation

Soil samples were collected during the 53^rd^ and 55^th^ Chilean Antarctic Scientific Expedition, during January-February 2017 and 2019. Samples were collected in triplicates, removing a 10 cm surface layer and then introducing ∼500 g of the underlying soil into Whirl-Pak sterile bags (Sigma) using an aseptic spade. Sampled zones covered the following islands: King George, Greenwich, Barrientos, Doumer, Deception, and Robert I, and included humanized areas (Gabriel de Castilla Base, Luis Risopatron Refugee, Yelcho Base, Henryk Arctowski Base, and Arturo Prat Base), as well as areas devoid of evident human intervention and restricted access (Fumarola Bay, Coppermine Peninsula, South Bay, Ecology Glacier, Air Force Glacier, and Barrientos Island). Further details of the sampled zones are provided in Supplementary Table 1. Soil bacteria were isolated by mixing 10 g of soil with 10-15 mL sterile PBS buffer, shaking in vortex for 5 min, and then let stand to allow the sedimentation of large solids. Then, 1 ml of the supernatant was put into a fresh tube and centrifuged at 1500 rpm for 1 min, plating 100 µl of the supernatant on nutritive agar plates supplemented with 100 µg/mL Cycloheximide. We used different culture media to increase the recovered diversity, including Marine agar, Casein Starch agar, Luria-Bertani (LB) agar, Tryptic Soy agar, Streptomyces Agar, and Actinomycete agar. Upon inoculation, the plates were incubated at 10-15ºC for approximately three weeks or until noticeable colonies formed. The obtained colonies were named and classified according to the site of sampling, Gram-staining results, and the morphology of the colonies and cells. For long-term storage of the Antarctic isolates, saturated liquid cultures were mixed 1:1 with glycerol 50% and kept at -80ºC.

### Antimicrobial sensitivity testing

To carry out antibiotic sensitivity tests with Antarctic bacteria, we adapted the disk diffusion method recommended by the European Committee for testing Antimicrobial Susceptibility (EUCAST, 2017). First, cultures of the strains to be tested were prepared in 2 mL of Mueller-Hinton broth (MH) and grown at 10° C with shaking for 48 h or until reaching the desired turbidity. Subsequently, the OD^625 nm^ of each culture was adjusted to 0.3 diluting with the same MH medium to obtain 2 mL of the adjusted culture. The adjusted culture was torula plated on MH agar to form a lawn. Subsequently, antibiotics discs (Oxoid) were applied on the inoculated plates using a disk dispenser (Oxoid). The plates were incubated at 10 °C for one week, after which the presence of growth inhibition halo was evaluated. The following disks were used: Amikacin 30 µg, Cyprofloxacin 5 µg, Ceftaroline 5 µg, Colistin 10 µg, Clindamycin 2 µg, Erythromycin 15 µg, Cefepime 30 µg, Fosfomycin/Trometramol 200 µg, Linezolid 30 µg, Meropenem 10 µg, Rifampicin 5 µg, Trimethoprim/sulfamethoxazole 25 µg, Tigecycline 15 µg, Piperacillin/Tazobactam 110 µg, and Vancomycin 30 µg. A strain was considered sensitive when a growth inhibition halo with a diameter equal to or greater than 10 mm was recorded.

### 16S rRNA-based taxonomic identification of isolates

Taxonomic classification of selected isolates was performed by PCR amplifying the full-lenght 16S rRNA gene using the primers 16SF (5’AGA GTT TGA TCC TGG CTC AG) and 16SR (5’ ACG GCT ACC TTG TTA CGA CTT). The purified amplicons were subjected to Sanger sequencing hiring the services of Macrogen Inc. (Korea) and the resulting sequence was compared to the NCBI 16S rRNA sequence database.

### Whole genome sequencing and analysis of Antarctic Isolates

Genomic DNA was extracted from selected isolates using the GenJet genomic DNA purification kit (ThermoFisher), following the manufacturer’s manual. DNA integrity and purity was asessed by agarose gel electrophoresis and measuring the ratio of absorbance at 260 and 280 nm. Prior to sequencing, DNA was quantified using a Qubit fluorimeter (ThermoFisher). To obtain complete genomes with closed chromosome and plasmids, we sequenced each genome using both illumina and Oxford Nanopore technologies. illumina sequencing was performed in a HiSeq 2500 platform using the Truseq DNA PCR Free library Kit (350 bp), at >50X coverage (Macrogen Inc., Korea). Nanopore sequencing was performed at Laboratorio de Expresión Génica (INTA, Santiago, Chile) using a MinION device (Nanopore, Oxford, UK) and with FLO-MIN106 (R9.4) flow cells, according to the manufacturer’s instructions. Sequencing libraries were prepared from 1 µg of gDNA using the 1D Genomic DNA by ligation kit (SQK-LSK108). Base-calling was performed using Guppy software (Nanopore, Oxford, UK). Genome assembly was performed using CANU (Koren et al., 2017) and Unicycler (Wick et al., 2017). The quality of the illumina reads was assessed using FASTQC (Andrews et al., 2012), and filtered using Trimommatic (Bolger et al., 2014). Adaptor sequences were removed from Nanopore reads using the tool Porechop (https://github.com/rrwick/Porechop). Genome annotation was performed using Prokka (Seemann, 2014). Antibiotic resistance gene identification was performed using the RGI-tool of the Comprehensive Antibiotic Resistance Database (Jia et al., 2017). Genome-based taxonomic assignment was performed using the rMLST (Ribosomal Multilocus Sequence Typing) analysis tool (Jolley et al., 2012) available at PubMLST.org (Jolley & Maiden, 2010) and the GTDB-Tk tool (Chaumeil et al., 2019). The phylogenomic tree showing the relationships between the Antarctic multiresistant isolates ArH3a and YeP6b, with other *Pseudomonas* was built calculating the Mash distances among the whole chromosomes Mash distances using Mashtree (Katz et al., 2019) with 1,000 bootstrap iterations.

### DNA extraction from Antarctic soil samples

Metagenomic DNA from soil samples was extracted using the DNeasy PowerSoil DNA isolation Kit (Qiagen), starting from 0.5-1 g of soil and following the manufacturer’s guidelines. Alternatively, soil DNA was extracted using a custom protocol. Briefly, 10 g of soil was mixed with 50 mL of Soil Lysis Buffer pre-heated to 65ºC (7.1 g Na_2_HPO_4_, 43.8 g NaCl, 5 g CTAB, 50 mL 1M Tris pH=8.0, 100 mL 0.5 M EDTA, nanopure water to complete 1 L), and Proteinase K. The mix was vortexed 30s and incubated at 60ºC for 2 h, mixing in vortex every 30 min. Then, 200 µg RNAse A were added and the mix was incubated at room temperature during 45 min shaking in vortex every 15 min, then centrifuging at 4000 x g during 2 min. The supernatant was transferred to a clean tube and then centrifuged 20 min at 11.000 x g. The supernatant was transferred to a clean tube and mixed with 1 vol. of 100% ethanol. The mixture was then sucesively loaded into a microfuge silica column, centrifuging during 2 min at 11000 x g and discarding the flow-through. Then, the DNA was eluted from the column using 100 µL of nanopure water, and then further purified using the reagents included in the DNeasy PowerSoil kit, mixing the DNA with 300 µL of the PowerBead buffer and performing the following steps recommended by the manufacturer. Metagenomic DNA integrity and purity was asessed by agarose gel electrophoresis and measuring the ratio of absorbance at 260 and 280 nm, respectively. Prior to sequencing, metagenomic DNA samples were quantified using a Qubit fluorimeter (ThermoFisher).

### 16S-rRNA amplicon sequencing and microbial diversity analyses

16S amplicon libraries were constructed and sequenced hiring the services of Macrogen Inc. (Korea). Upon quality control, metagenomic DNA extracted from soil samples was used along with the Herculase II Fusion DNA Polymerase Nextera XT Index Kit V2 (illumina), in order to amplify the V3-V4 region of the 16S rRNA gene using the primersBakt_341F: CCTACGGGNGGCWGCAG and Bakt_805R: GACTACHVGGGTATCTAATCC. The libraries were sequenced using an illumina Miseq sequencer, obtaining approximately 166,000-245,000 300 bp paired-end reads (50-73 Mbp) per sample. Microbial diversity analyses including relative abundance, and alpha- and beta-diversity calculations were performed using QIIME2 platform v. 2019.7.0 (Bolyen et al., 2019). The 16S rRNA amplicon reads were imported into the Qiime2 pipeline, performing quality assessment and filtering using q2-dada2. The result obtained was explored using featureTable function. A rooted phylogenetic tree required for alpha and beta diversity analyses was generated with the q2 -diversity plugin, performing PERMANOVA statistical analysis for beta diversity. On the other hand, the compositional analysis of beta diversity was carried out using the DEICODE complement. Finally, q2-feature-classifier was used to carry out the taxonomic assignment of the microorganisms present in the samples using the SILVA 138 database.

### Shotgun metagenomic sequencing

For illumina shotgun metagenomic sequencing of samples collected at A. Prat base, Air Force glacier, Gabriel de Castilla base, and Barrientos island (during ECA53), the libraries were prepared from soil DNA using the Truseq DNA PCR Free library Kit (350 bp) and were sequenced in an illumina Hiseq2500 platform, obtaining a total of 4.9 to 6.7 Gbp per sample (100 bp paired-end reads). Additionally, the libraries from samples collected at H. Arctowski Base, Coppermine Peninsula, and Fumarola Bay (during ECA55) were prepared using the Truseq nano DNA library Kit (350bp), and were sequenced in an illumina Novaseq sequencer, obtaining a total of 11.8-12.7 Gbp per sample (150 bp paired-end reads). Nanopore sequencing libraries were prepared starting with 1 µg of soil DNA using the Ligation Sequencing kit LSK-109, following the manufacturer’s guidelines. For each library, 5-50 fmol were loaded into a R9 flow cell and sequenced using a MinION device (Nanopore Technologies).

### Metagenome assembly

Prior to assembly, illumina reads were quality-assessed using FastQC and trimmed (adaptors and low quality bases) using Trimmomatic (Bolger et al., 2014). The quality of Nanopore reads was evaluated using NanoPlot, while adaptor removal was performed using Porechop (https://github.com/rrwick/Porechop). Hybrid metagenome assembly was performed combining pre-processed illumina and Nanopore data using metaSPAdes (Nurk et al., 2017). The quality of the obtained assemblies was assessed using metaQUAST (Mikheenko et al., 2016).

### ARGs relative abundance calculations based on illumina reads

Antibiotic resistance gene identification starting from unassembled illumina data was performed using the tool deepARG (Arango-Argoty et al., 2018), recovering the “mapping.ARG” output file, which contains alignments that have a probability higher or equal to 0.8, which according to the authors, is reliable enough to be considered as a true positive alignment. To construct the sunburst chart showing the most abundant ARG in the sum of the sampled sites, all the ARG alignments from all sites were added in a single table. Then, ARGs with less than 1000 alignments were filtered out, and the number of alignments for each surviving ARG was normalized by the total number of alignments for all of the ARGs. For relative abundance normalization among different samples, the illumina reads from each metagenome were mapped to a custom *rpoB* protein sequence database (kindly provided by Dr. Luis Orellana) using DIAMOND BlastX, with cutoffs of 80% identity and 1E-10 e-value for positive alignments. ARG abundances for each metagenome were calculated considering the total amount of *rpoB* alignments as a normalizing factor, the known length of each ARG in the deepARG database, and the median length for *rpoB* sequences (4029 bp). The deepARG-abundance module from the genetools software was used for this task. For constructing the heatmaps showing the ARGs relative abundance among different sites, seriation distances were determined using the GW method and the seriation R library. The final Heatmap vector graphics were created using the ComplexHeatmap and circlize R libraries. Discriminant ARGs analysis was performed using the Extrarg tool, using as input a table comprising the known amount of raw ARG alignments in every metagenome and the corresponding normalized amount. This table also included a group designation (humanized or non-intervened). ARG categorization by resistance mechanism, drug class, and ARG gene family was performed by searching in the CARD database each of the matched genes from deepARG analysis. NMDS plots were created using the Vegan and ggplot2 R packages. Permanova analysis was performed using the pairwiseAdonis R package.

### ARGs identification in metagenomic assemblies and mobile genetic elements prediction

ARG identification among assembled metagenomic contigs was performed using NanoARG (Arango-Argoty et al., 2019). Genetools was used to convert the .json output file into a table that includes information of all the hits (i.e., identity, coverage, e-value). To construct the treemap showing the most abundant ARGs detected in the sum of the assemblies, we concatenated the output tables obtained for each sample. Next, we examined the distribution of the percentage identity and e-value among all the hits to establish appropriate cutoffs, which allow minimizing false positives but permitting the detection of novel resistance genes with limited identity to those present in the database. Based on the observed data distribution, we established a 40% identity and 1E-10 e-value cutoff. The ARG relative abundance was calculated by dividing the number of hits obtained for each ARG by the total number of hits for all the ARGs. Taxonomic assignment of contigs carrying ARGs was performed using Centrifuge (Kim et al., 2016). Plasmid prediction among assembled metagenomic contigs was conducted using PlasFlow (Krawczyk et al., 2018) and ViralVerify (Antipov et al., 2020).

### Metagenome-assembled genomes (MAGs) isolation and characterization

MAGs recovery was performed as follows. The hybrid metagenomic assembly generated using metaSPAdes, as well as the short and long read sets from each metagenome, were used as input for MetaWRAP (Uritskiy et al., 2018) in order to perform contig binning and initial bins refinement using a combination of MetaBAT2 (Kang et al., 2019), CONCOCT (Alneberg et al., 2014), and MaxBin2 (Wu et al., 2016) binning algorithms. Afterward, the contigs from the obtained bins were used as a reference to map the whole set of long and short reads using minimap2 v2.17 (Li, 2018) and Bowtie2 (Langmead & Salzberg, 2012), respectively. Upon recovering, the mapping reads were combined, and de novo assembled using Unicycler. The completeness and contamination of the MAGs were evaluated using CheckM (Parks et al., 2015). MAG taxonomic assignment was conducted using GTDB-Tk. In the case of MAG 22 from Air Force Glacier, GTDB-Tk predicted it as Dormibacteriota, although analysis based on genes coding for 50S ribosomal proteins L6, L14, L16, L24; 30S ribosomal proteins S1, S5, S8, S9, S11; and Elongation factors Tu and G, indicated that this MAG would belong to Phylum Chloroflexota as well as the rest of the members of the same clade, as shown in the phylogenetic tree of Figure 8. This MAGs tree was constructed using the SpeciesTreeBuilder v.0.1.0 from the Kbase App catalog (Arkin et al., 2018), based on the alignment of 49 core universal COG (Clusters of Orthologous Groups) gene families. The obtained tree was formatted using FigTree v1.4.4 (https://github.com/rambaut/figtree/releases). MAGs general annotation was performed using Prokka. ARGs identification in MAGs was performed using RGI-CARD tool, allowing loose hits.

### Antarctic OXA beta-lactamase structure modeling

For homology-based structural modeling of the metagenomic OXA-like beta-lactamase, the amino acid sequence was used to search the Uniprot database for proteins with 3D structure using BLAST on the online server. Three templates were chosen with an identity percentage greater than 50%, corresponding to class D beta-lactamases identified in *Klebsiella pneumoniae* (PDB 6XJ3), *Salmonella enterica* subsp. enterica serovar Typhimurium (PDB 1K38) and *Pseudomonas aeruginosa* (PDB 3IF6). The templates and the target sequence were aligned, and 50 models were built using Modeller v10.1 (Šali & Blundell, 1993). Then, we evaluated the models considering the results of the DOPE score (Shen & Sali, 2006), QMEAN (Benkert et al., 2011), and MolProbity (Williams et al., 2018), selecting the best structure from these evaluations. The selected structure was visualized and formatted in PyMOL v2.4 (L DeLano, 2002), highlighting the relevant residues from the active site, as described in (Paetzel et al., 2000).

## RESULTS AND DISCUSSION

### Antibiotic-resistant bacteria in Antarctic soil

We examined the antibiotic resistance capabilities of 229 psychrotolerant bacterial strains isolated from soil collected from different locations of North Antarctica during the 53_rd_ and 55_th_ Chilean Scientific Antarctic Expedition (January-February 2017 and 2019, respectively), covering areas intervened by human activities (i.e., the surroundings of human bases) as well as restricted areas devoid of evident human intervention (Figure 1A, Supplementary Table 1). We tested the isolates’ sensitivity to fifteen clinically used antibiotics of diverse chemical nature and mechanism of action using adapted disk diffusion assays. We found at least one resistant isolate to 13 out of 15 antibiotics tested, while only meropenem and tigecycline could kill all the isolates (Figure 1B). In addition, a high proportion of resistant isolates was observed for Fosfomycin and Colistin, followed by Clindamycin and Vancomycin.

**Figure 1.**
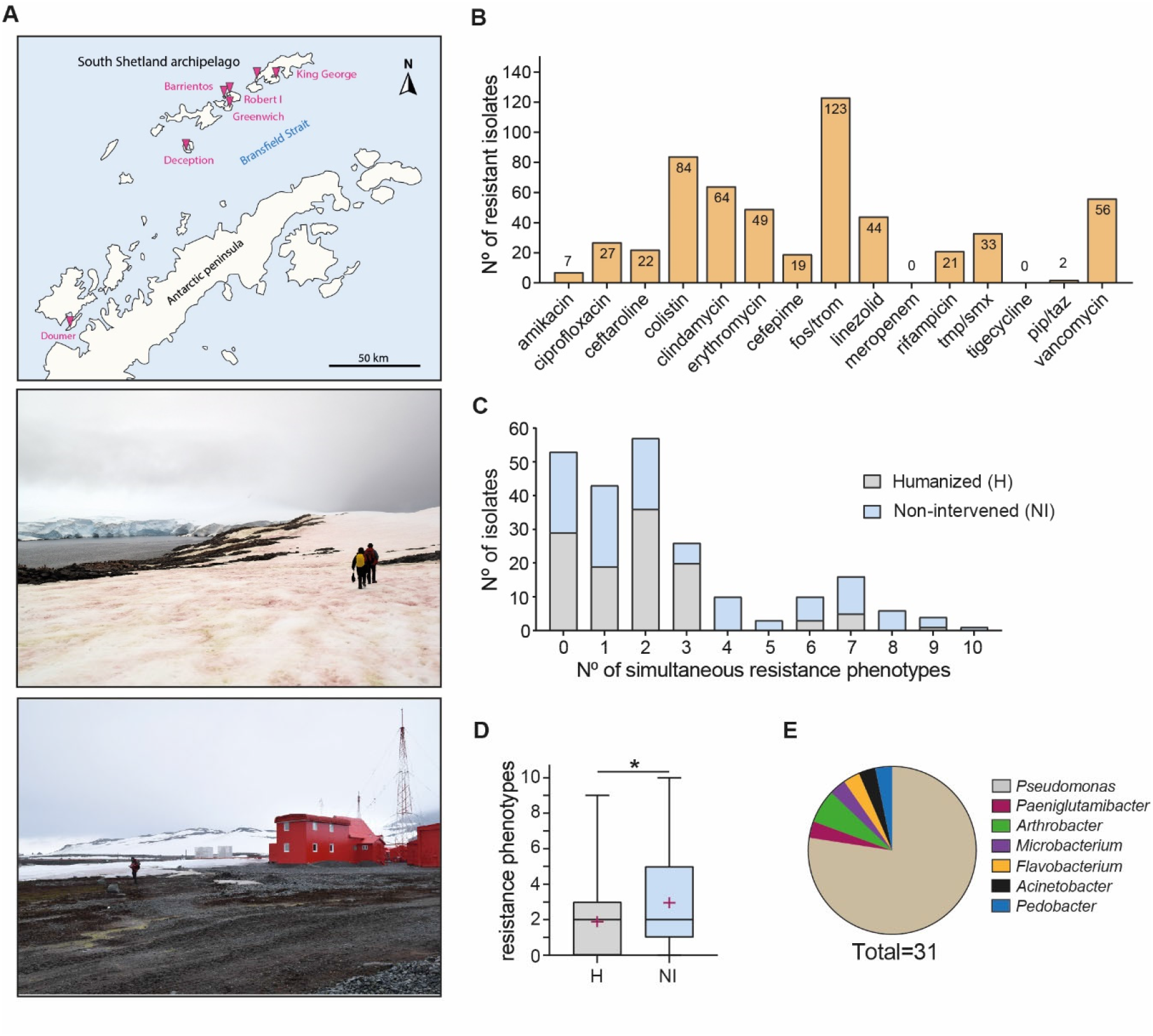
Isolation of antibiotic resistant bacteria from North Antarctica soils. (A) Soil samples were collected at different locations of North Antarctica (pink triangles), including areas devoid of noticeable human intervention (upper picture) and areas harbouring human settlements (lower picture). (B) Antibiotic sensitivity testing of Antarctic bacterial iso lates using adapted disk fifussion assays. (C) Recount of isolates showing simultaneous resistance to one or more antibiotics, both from humanized or non-intervened areas. (D) Average number of simpultaneos resistances displayed by Antarctic isolates coming either from humanized or non-intervened areas. (E) Taxonomic classification of selected Antarctic isolates showing resistance to 7 or more antibioti cs, or showing resistance to antibiotics that killed most of the isolates (e.g. piperacillin/tazobactam).

Next, we assessed if the resistance to each antibiotic could be preferentially associated with isolates obtained from either humanized or non-intervened areas, looking for possible enrichment of resistant bacteria due to anthropogenic activities (Supplementary Table 2). We found significant associations for eight of the fifteen antibiotics tested (odds ratio: 0.10-5.87; p<0.05, two-tailed Fischer’s exact test) (Table 1). In seven of them, the resistance had three to six-fold increased probability among non-intervened areas, arguing against increased resistance among soil bacteria due to anthropogenic intervention. The only exception was ciprofloxacin, with a 10-fold increased probability of resistance in humanized areas. This is in agreement with a previous report showing the presence of ciprofloxacin in wastewater from different humanized areas in King George Island (Hernández et al., 2019).

**Table 1.** Comparison of antibiotic resistance among soil bacteria isolated from either humanized or non-intervened areas from north Antarctica.

| Antibiotic | Non-intervened<br>(n=116) |  | Humanized<br>(n=113) |  | Statistics<sup>1</sup> |  |  |
| --- | --- | --- | --- | --- | --- | --- | --- |
|  | n | % | n | % | p-value | OR | 95% CI |
| Amikacin | 6 | 5.2 | 1 | 0.9 | 0.1195 | - | - |
| Ciprofloxacin | 3 | 2.6 | 24 | 21.2 | <0.0001 | **0.10** | 0.03 - 0.33 |
| Ceftaroline | 17 | 14.7 | 5 | 4.4 | 0.0123 | **3.71** | 1.32 - 9.42 |
| Colistin | 38 | 32.8 | 46 | 40.7 | 0.2205 | - | - |
| Clindamycin | 45 | 38.8 | 19 | 16.8 | 0.0002 | **3.14** | 1.70 - 5.66 |
| Erythromycin | 39 | 33.6 | 10 | 8.8 | <0.0001 | **5.22** | 2.41 - 10.54 |
| Cefepime | 16 | 13.8 | 3 | 2.7 | 0.0031 | **5.87** | 1.76 - 19.35 |
| Fosfomycin/<br>trometamol | 60 | 51.7 | 63 | 55.8 | 0.5966 | - | - |
| Linezolid | 34 | 29.3 | 10 | 8.8 | <0.0001 | **4.27** | 2.05 - 8.69 |
| Meropenem | 0 | 0.0 | 0 | 0.0 | - | - | - |
| Rifampicin | 15 | 12.9 | 6 | 5.3 | 0.0654 | - | - |
| Trimethoprim/<br>sulfamethoxazol | 24 | 20.7 | 9 | 8.0 | 0.0079 | **3.01** | 1.39 - 7.03 |
| Tigecycline | 0 | 0.0 | 0 | 0.0 | - | - | - |
| Piperacillin/<br>tazobactam | 2 | 1.7 | 0 | 0.0 | 0.4979 | - | - |
| Vancomycin | 42 | 36.2 | 14 | 12.4 | <0.0001 | **4.01** | 2.09 - 7.89 |

<sup>1</sup>p-value calculated using a two-tailed Fisher's exact test; OR: odds ratio calculated for significant differences (p<0.05); CI: OR confidence interval calculated following the Baptista-Pike method.

Following, we evaluated the occurrence of multi-resistance, identifying isolates resisting up to ten different antibiotics (Figure 1C). From the total isolates, 50 (∼22%) were resistant to four or more of these compounds, of which 41 came from non-intervened areas. Conversely, most of the isolates from humanized areas showed resistance to three or fewer antibiotics. In this direction, isolates from non-intervened areas averaged a significantly higher number of simultaneous resistance phenotypes (p<0.05, two-tailed Mann-Whitney test) (Figure 1D). Twenty-seven isolates showing resistance to seven or more antibiotics were classified taxonomically based on 16S-rRNA analysis. Twenty-four of them were assigned to the genus *Pseudomona*s (Proteobacteria), one to *Pedobacter* (Bacteroidetes), and one to *Paeniglutamicibacter* (Actinobacteria) (Figure 1E). On the other hand, isolates showing resistance to amikacin or tazobactam/piperacillin (antibiotics with the lowest proportion of resistance among the whole collection) were classified as *Microbacterium* (Actinobacteria), *Acinetobacter* (Proteobacteria), *Flavobacterium* (Bacteroidetes), and *Arthrobacter* (Actinobacteria). These results indicate a high proportion of antibiotic-resistant isolates in the sampled Antarctic soils, with no evidence of anthropogenic-derived enrichment. Most of the multiresistant isolates came from non-intervened areas and corresponded to *Pseudomonas*.

### Genomic features of multiresistant Antarctic isolates

To investigate possible genetic determinants behind the multi-resistance phenotype, we selected two Antarctic isolates as models for genomic analyses: *Pseudomonas* YeP6b and ArH3a, showing resistance to 10 and 9 antibiotics, respectively (Figure 2A). Using both Illumina and Nanopore sequencing, we generated ∼5.7 Gbp short reads (150 bp, paired-end) and 800 Mbp long reads for YeP6b, and ∼6.7 Gbp short reads plus ∼5.7 Gbp long reads for ArH3a, which were combined in each case to build hybrid assemblies. This way, YeP6b showed a ∼6.66 Mbp closed chromosome with no detected plasmids, while ArH3a showed a ∼6.77 Mbp closed chromosome and a 2,628 bp small plasmid (pArH3a). Although coming from roughly 420 km far apart (Doumer and King George islands), both strains shared 99.46% average nucleotide identity (ANI), indicating they are members of the same species. Genome-based taxonomic placement using GTDB-Tk (Chaumeil et al., 2019) indicated *Pseudomonas* sp. IB20 (GCF_002263605.1) as the closest genome for both YeP6b and ArH3a, showing an ANI value of 99.28% and 99.36%, respectively. *P*. sp. IB20 was previously isolated from King George Island and classified inside the *Pseudomonas fluorescens* species complex, within the *P. antarctica* group and distant from animals or plants pathogens such as *P. aeruginosa* and *P. syringae* (Vásquez-Ponce et al., 2018). Phylogenomic analysis using Mashtree (Katz et al., 2019) supported this grouping (Supplementary Figure 1, Supplementary Table 3). Thus, the multiresistant isolates YeP6b and ArH3a are part of a ubiquitous bacterial species naturally inhabiting North Antarctica’s soils.

**Figure 2.**
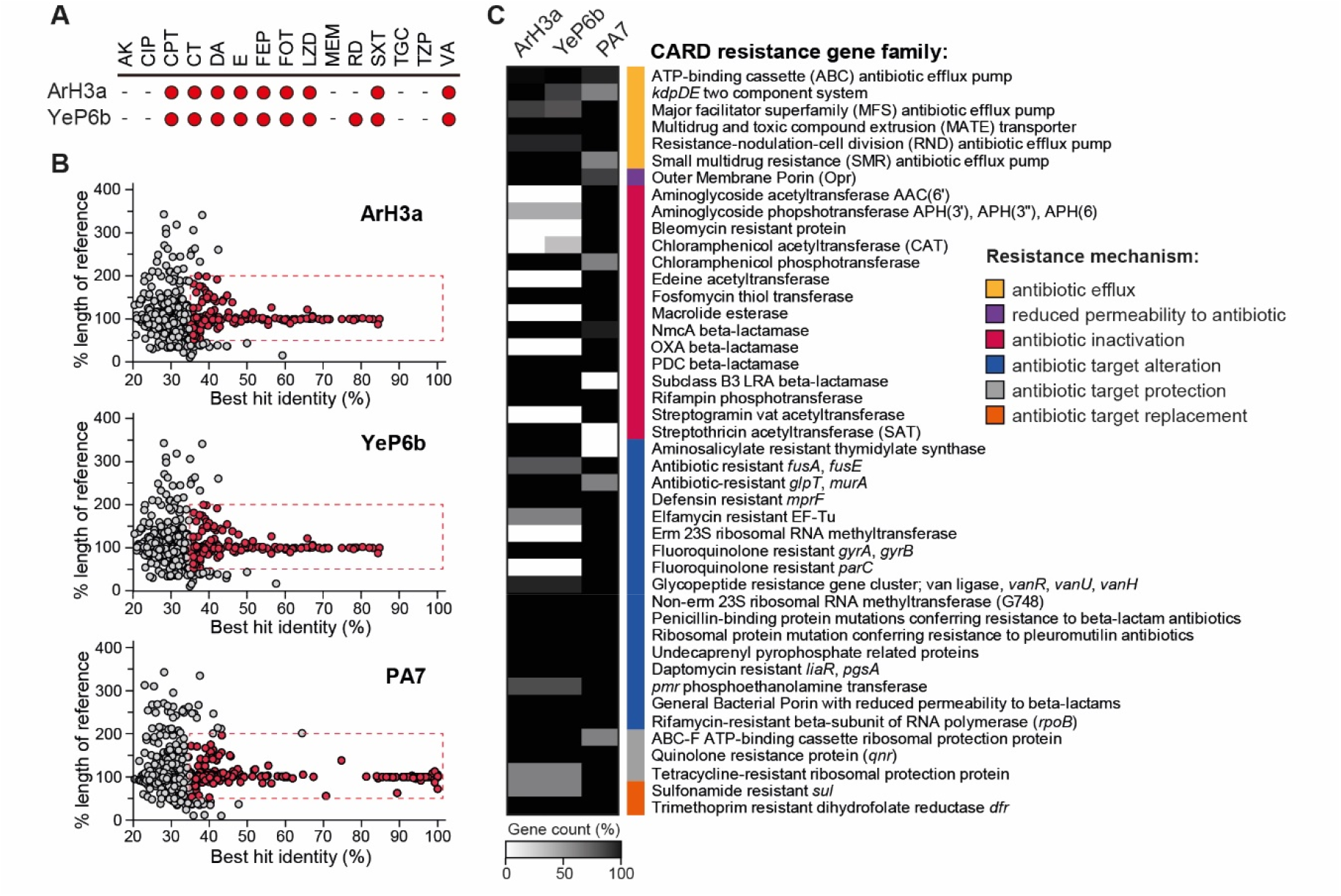
Experimental and genomic antibiotic resistance profile of the multirresistant Antarctic isolates *Pseudomonas* spp. ArH3A and YeP6b. (A) Experimental sensitivity to antibiotics determined for each Antarctic isolate. (B) Distribution of the percentage identity and the reference coverage of hits obtained upon searching for ARGs in the indicated genomes using RGI-CARD . The hits showed in red were selected to construct the heatmap shown in (C). (C) Heatmap showing the abundance of ARG gene families from different classes among Antarctic *Pseudomonas* ArH3a and Yep6b, and from the multiresistant clinical isolate *Pseudomonas aeruginosa* PA7. The gene count was calculated by dividing he number of all the detected ARGs falling in the mentioned category in each genome, divided by the highest number of ARGs.

Genome annotation indicated 6,338 protein-coding genes in the *Pseudomonas* sp. ArH3a chromosome, among other features (Supplementary Table 4). Additionally, pArH3a had 5 CDS encoding a putative plasmid replication protein and four hypothetical proteins without predicted function. Likewise, we identified 6,141 CDS in the *Pseudomonas* YeP6b chromosome. Prophage content analysis using Phaster (Arndt et al., 2016) revealed eight of them in the chromosome of ArH3a and seven in YeP6b, sharing only one of them, reflecting intraspecies genome diversity due to horizontal gene transfer. Next, we used the tool Resistance Gene Identifier (RGI) from the Comprehensive Antibiotic Resistance Database (CARD) to identify and classify putative antibiotic resistance genes (ARGs) present in these genomes. For comparison, we included in the analysis the genome of the virulent clinical isolate *Pseudomonas aeruginosa* PA7, previously studied as a model multiresistant strain (Roy et al., 2010; Singh et al., 2020). RGI allows ARG identification under three paradigms: perfect, strict, or loose. The Perfect and Strict algorithms detect perfect matches to the curated reference sequences or previously unknown variants of known AMR genes and are best suited for clinical surveillance of known bacterial pathogens (Jia et al., 2017). We found no perfect hits and only four strict hits for both YeP6b and ArH3a, including a homolog of the *P. aeruginosa* transcriptional activator SoxR linked to increased expression of RND efflux pumps (Palma et al., 2005; Sporer et al., 2018). Also, two genes similar to the AdeF RND efflux pump and one homolog to the AbaQ MFS efflux pump, both of them described in *Acinetobacter baumannii* and reported to confer resistance to tetracycline, fluoroquinolones, and other antibiotics (Coyne et al., 2010; Pérez-Varela et al., 2018). RGI analysis indicated five perfect and forty-one strict hits for PA7, including genes encoding four aminoglycoside phosphotransferases and one acetyltransferase, the Cmx chloramphenicol exporter, an OXA-50 beta-lactamase, and several efflux pumps (i.e. MexAB-OprM, MexCD-OprJ, and MexXY). The lower number of ARGs found in the multiresistant Antarctic isolates suggests that functionally similar genes sharing limited identity with known ARGs or genes encoding different molecular mechanisms would explain their resistance phenotype.

To get a more comprehensive view of possible genes involved in Antarctic isolate multi-resistance, we included in the analysis the loose hits, recommended for new resistance genes discovery in bacteria different from clinical isolates, although with increased expected false positives (Jia et al., 2017). We found 700 and 701 additional putative ARGs in YeP6b and ArH3a, respectively, and 671 in PA7. As a way to identify and filter out possible false positives, we plotted for each hit the percentage identity and length to the reference ARG, noticing a high dispersion of the coverage length below 35% identity (Figure 2B). In contrast, we found a clear tendency to similar length at increasing identity values. Thus, to focus on the genes more likely to share functions with known resistance genes, we decided to include the hits showing ≥35%+ identity and 50-200% length of the reference. This way, 168, 165, and 228 hits remained for ArH3a, YeP6b, and PA7, respectively. We grouped all the putative ARGs by gene family and resistance mechanism, then calculating each category’s relative gene count in each strain (Figure 2C). Antarctic isolates showed an overall lower number of predicted ARGs than *P. aeruginosa* PA7, lacking genes encoding aminoglycoside acetyltransferase AAC(6’), macrolide esterases, OXA beta-lactamases, streptogramin vat acetyltransferases, and Fluoroquinolone-resistant parC. However, the Antarctic isolates showed a higher number of genes encoding ATP-binding cassette (ABC) antibiotic efflux pumps and small multidrug resistance (SMR) antibiotic efflux pumps than PA7. Also, ArH3a and YeP6b strains shared with PA7 a putative carbapenem-hydrolyzing Class-A NmcA beta-lactamase originally described in *Enterobacter cloacae* (Naas & Nordmann, 1994), and a putative class-C PDC beta-lactamase, which is generally found in *Pseudomonas aeruginosa*, conferring resistance to monobactam, carbapenems, and cephalosporins (Rodríguez-Martínez et al., 2009). Interestingly, among the putative ARGs found in the Antarctic isolates but absent in PA7, we found a homolog of the LRA-3 beta-lactamase, previously characterized from metagenomic DNA from soil samples in Alaska. Notably, it was demonstrated that this beta-lactamase confers resistance to cephalosporins and penam antibiotics when expressed in *E. coli* (Allen et al., 2009).

All these results indicate that Antarctic soil culturable microbiota is a source of genes that, although encoding proteins sharing limited identity with known resistance genes, would retain the capability to confer resistance to various antibiotics. This microbiota would also host genes encoding novel resistance mechanisms that could be further studied to anticipate possible new resistance mechanisms arising in the future among bacterial pathogens.

### Microbial diversity in North Antarctica soils

To get a detailed view of the microbial diversity inhabiting the sampled soils and compare the communities found in humanized and non-intervened areas, we performed 16S rRNA sequencing-based analyses. Relative abundance calculations indicated that Proteobacteria, Bacteroidota, Acidobacteriota, and Verrucomicrobiota corresponded to the most abundant Phyla in most soils (Figure 3A), while among the most abundant genera were *Candidatus Udaeobacter*, RB41, *Polaromonas*, and *Ferruginibacter* (Figure 3B). Accordingly, a recent study of microbial diversity in recently exposed soils from the Midtre Lovénbreen glacier in the Arctic with similar weather conditions that North Antarctica also reported a high proportion of *Candidatus Udaeobacter* (Verrucomicrobiota) and RB41 (Acidobacteriota), uncultured Gemmatimonadota and *Polaromonas* (Venkatachalam et al., 2021). This is also consistent with a previous study correlating the air temperature with microbial diversity in maritime Antarctica soils, showing *Polaromonas* and *Gemmatimonas* among the dominant taxa (Dennis et al., 2019). Of note, a marked decrease in *Candidatus Udaeobacter* and RB41 was observed in Antarctic humanized soils, compared to non-intervened areas (Figure 3B). These two genera were also reported to be enriched in Arctic soils in an older deglaciation stage (Venkatachalam et al., 2021). This suggests that these genera could be markers of recently exposed or non-intervened soils in polar habitats. Alpha diversity calculations (Shannon and Pielou) indicated a high microbial diversity in all the samples, showing no significant differences in average when comparing humanized vs. non-intervened sites (Figure 3C). On the other hand, beta diversity calculations based on Bray-Curtis dissimilarity and Robust Aitchison compositional analysis revealed differences in the composition of the microbial communities at each site, with statistically supported clustering of non-intervened sites apart from humanized areas (P=0.004 and 0.006, for Bray-Curtis and Aitchison, respectively) (Figure 3D). In sum, our analyses indicated that a highly diverse microbiota inhabit the sampled soils with a dominance of taxa also found in other polar environments. Furthermore, although microbial communities from non-intervened and humanized areas showed different compositions, this could not be directly attributed to non-indigenous taxa present in the latter.

**Figure 3.**
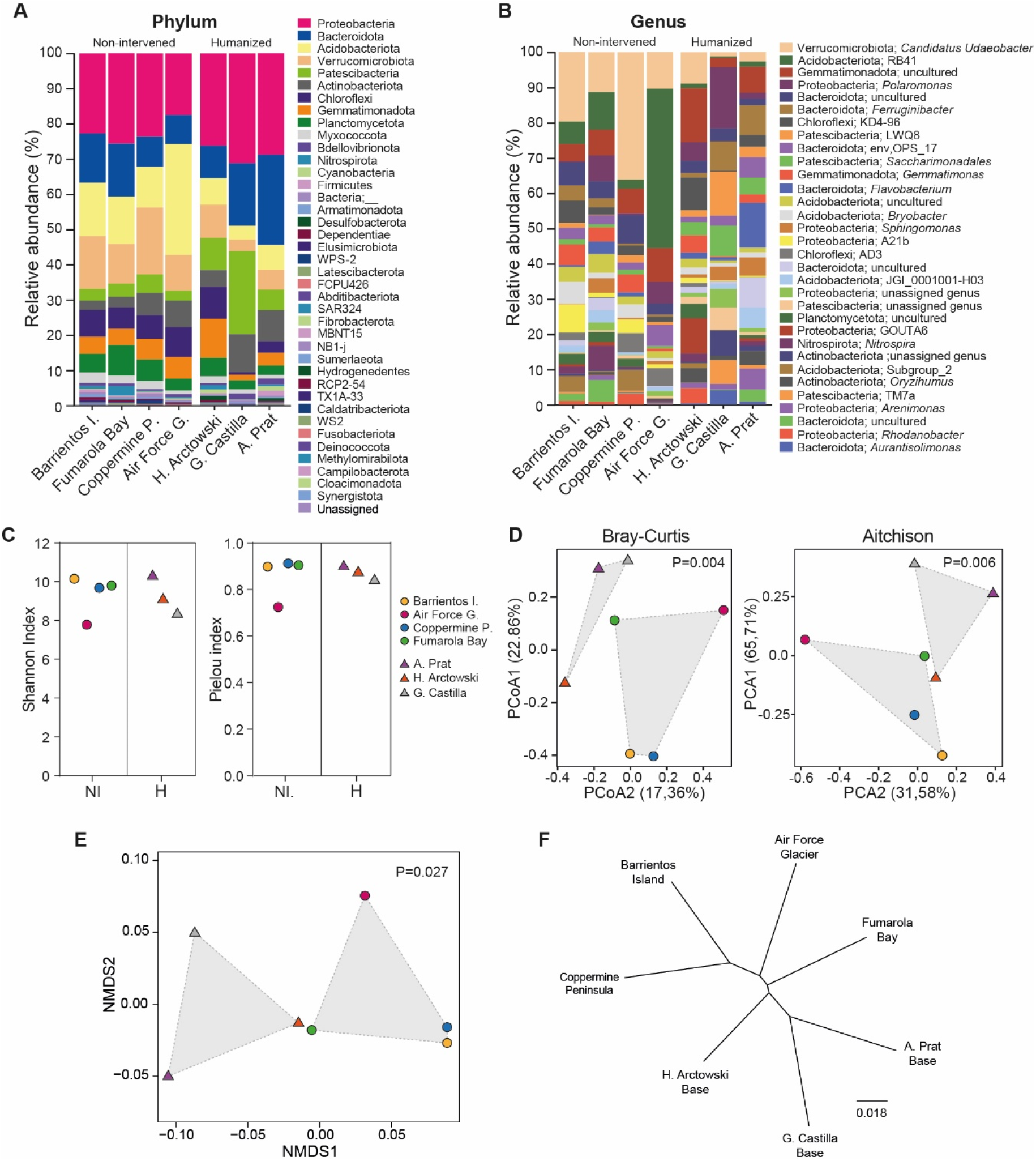
Microbial and DNA sequence diversity in the sampled Antarctic soils. (A-D) 16S rRNA-based microbial diversity analysis. Relative abundance of bacterial phyla (A) and genera (B) amongthe sampled sites. (C) Alpha diversity estimation based on Shannon and Pielou index. (D) Beta diversity estimation based on Bray-curtis dissimilarity index and the Robust Aitchison analysis. (E-F) Mash analysis of sequence diversity among the shotgun metagenomes of the analyzed soil samples. (E) Non-metric multidimensional scaling based on the Mash distances. Mashtree inferred from the Mash distances calculated for the set of metagenomes. The P-value supporting sample clustering as represented by gray polygons was calculated performing a PERMANO VA analysis.

### Detection of antibiotic resistance genes hosted by Antarctic soil bacteria

To evaluate the presence and abundance of ARGs in North Antarctica soils and investigating the microbiota hosting these factors, we conducted a set of shotgun sequencing-based metagenomic analyses. First, Illumina sequencing was performed for the metagenomes obtained from each of the four non-intervened and three humanized areas included in the microbial diversity assessments. In this regard, sequence diversity analysis using Mash corroborated the differential clustering of both kinds of areas (Figure 3E and 3F).

Illumina reads were used to conduct a mapping-based strategy to identify and calculate the relative abundance of ARGs using the tool deepARG and custom Python scripts. This approach has the advantage of being independent of metagenome assembly, which is highly dependent on sample sequence complexity and coverage (Rodriguez-R et al., 2018), hampering abundance comparisons between samples. However, as caveats, short read mapping does not allow the identification of complete ARGs and investigation of their genomic context. Overall, the mapped metagenomic reads from the sum of all the samples revealed the presence of hundreds of different antibiotic resistance gene families, which fall into more than 17 classes according to the drug to which they confer resistance (Figure 4A). As for the multiresistant Antarctic isolates, we found a high proportion (∼40%) of matches to genes encoding efflux pumps and ABC transporters that would confer resistance to multiple drugs. Furthermore, we found different genes coding for proteins potentially involved in antibiotic inactivation and target alteration, protection, or replacement, which would confer resistance to Macrolides, Streptogramin and Lincosamide (∼19%), tetracycline (∼10%), Bacitracin (∼7%), and glycopeptides (∼4%). Although with lesser abundance, we also found matches to genes conferring resistance to beta-lactams, aminoglycosides, rifamycin, fosmidomycin, and peptide antibiotics. These results agree with two previous studies of soil metagenomes from Mackay Glacier in South Victoria Land and King George Island in Antarctica, where a set of genes encoding putative efflux pumps and antibiotic inactivation enzymes were predicted (Goethem et al., 2018; Yuan et al., 2019).

**Figure 4.**
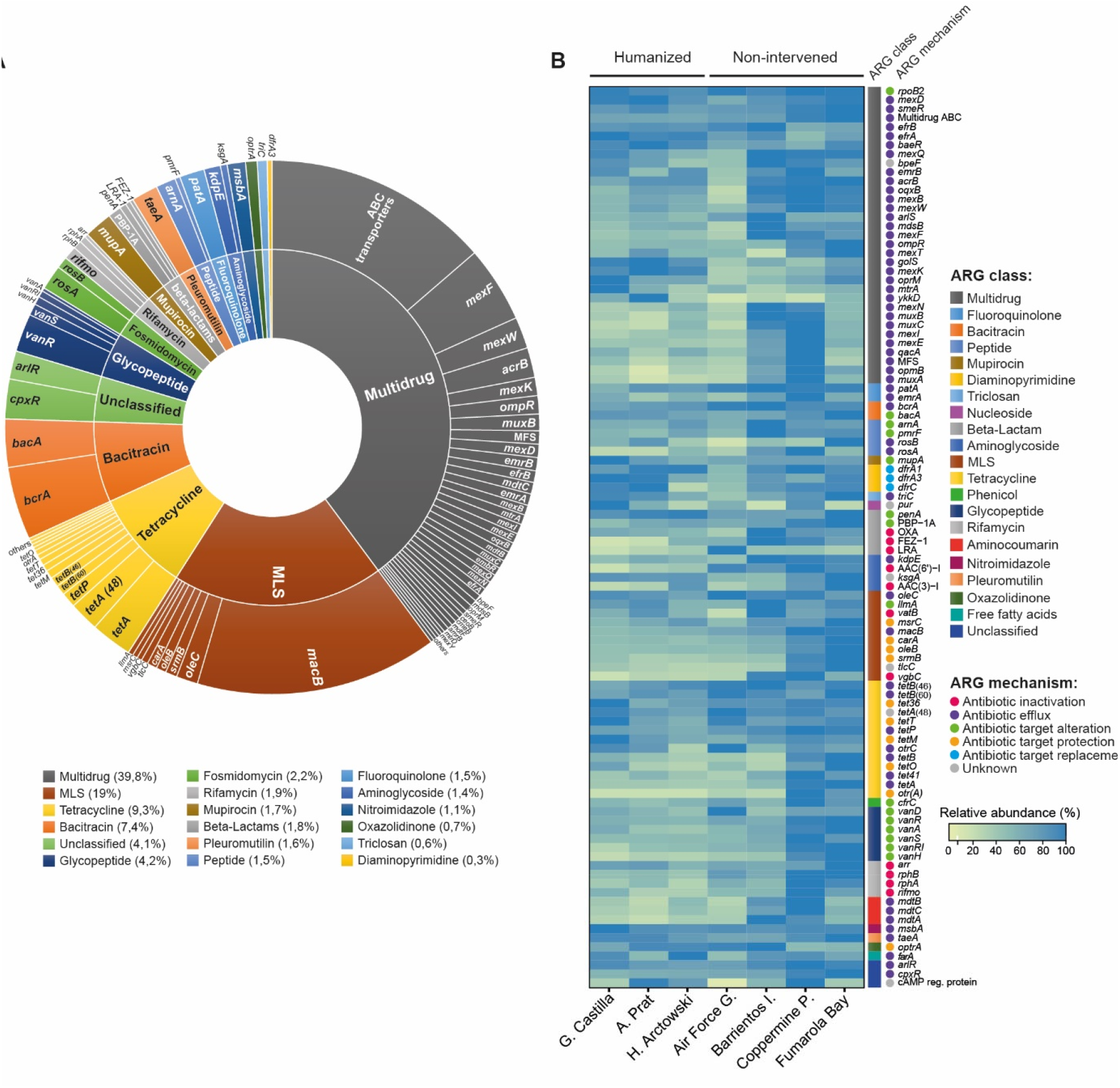
Variety and relative abundance of predicted ARGs among Antarctic soil microbiota based on short reads analysis. (A) Sunburst plot showing the proportion of hits found for each resistance gene family and drug class in the sum of the seven sampled sites. (B) Heatmap showing the normalized abundance of different ARG families in different Antarctic sites, calculated using a short-reads mapping based strategy. Normalization of the abundance comparing different sites was performed based on reads mapping to *rpoB*. Also, for each ARG the abundance was normalized by the highest value among the sites (for each column). For each ARG, the resistance gene family and drug class is indicated (based on CARD categorization).

To measure and compare the relative abundance of ARGs in different soils, we followed a normalization strategy based on using the *rpoB* gene as a single-copy universal bacterial marker to correct possible differences in total gene abundance across the sites. The relative calculated abundances were used to construct a heatmap including the 100 most abundant ARGs (Figure 4B). An overall higher abundance of different ARGs was noticed in three non-intervened sites (Barrientos Island, Coppermine Peninsula, and Fumarola Bay), especially for several efflux pumps and genes involved in resistance to MLS, glycopeptides, and rifamycin. Also, a widespread high abundance was observed for ABC transporters and several *tet* genes conferring resistance to tetracycline. To search for possible ARGs able to discriminate between soils from humanized and non-intervened areas, we used extrARG, a tool developed to identify discriminatory antibiotic resistance genes among environmental resistomes using an extremely randomized tree algorithm (Gupta et al., 2019). Twenty-three discriminatory ARGs were identified, covering different classes and mechanisms (Supplementary Figure 2A). The discriminatory power of these ARGs was assessed through NMDS analysis, obtaining a clear significant differential grouping of the samples (P=0.029) (Supplementary Figure 2B). Among the ARGs that were characteristic of humanized zones are *dfrK*, arr-5, *axyX, pmrE*, and the JOHN-1 beta lactamase. Conversely, ARGs characteristic of non-intervened zones are *cpxA*, AAC(6’)-I, *vgbC, mexM*, and *mel*.

To get a first approach of the microbial taxa harboring the ARGs that generated positive hits, we extracted all the reads mapping to any ARG and performed taxonomic assignment using Kaiju (Menzel et al., 2016). As a result, most of the reads mapping to different classes of ARGs were assigned to *Pseudomonas, Streptomyces, Gemmatimonas, Paenibacillus*, and *Polaromonas* (Supplementary Figure 3), supporting that the ARGs found in the soil metagenomes came from bacteria naturally inhabiting Antarctic soils rather than from introduced non-indigenous bacteria.

### Presence of antibiotic resistance genes in assembled contigs and metagenome-assembled genomes (MAGs)

To get further evidence of the presence of ARGs among the Antarctic soil metagenomes and to gain more information regarding these genes and their genomic context, we performed Nanopore sequencing of three of the seven metagenomes analyzed in the previous section (G. de Castilla Base, A. Prat Base, and Air Force Glacier). In combination with short reads, long Nanopore reads data allowed us to construct hybrid metagenome assemblies, resulting in longer contigs and larger total assembly size than short reads-only assemblies. Upon assembly, the contigs were annotated and searched for the presence of ARGs using the platform nanoARG (Arango-Argoty et al., 2019). Positive hits were defined with a cutoff of 40% identity and 1E-10 e-value, based on the observed distribution of percentage identity and e-value, to detect novel resistance genes although minimizing the false positives. We identified 2,214 ARGs among the contigs from the three assembled metagenomes, corresponding to 191 different resistance gene families, closely resembling the array of ARGs found in the short reads-based analysis (Supplementary Figure 4A). Assembled contigs comprising ARGs were subjected to taxonomic assignment using Centrifuge (Kim et al., 2016), corroborating *Polaromonas, Pseudomonas, Streptomyces, Variovorax, Burkholderia, and Gemmatimonas* among the main host taxa for the identified ARGs (Supplementary Figure 4B), in agreement with short reads analysis.

To gain a further detailed view of the uncultured microbial diversity present in North Antarctica soils and evaluate the presence of ARGs in a more genome-specific way, we applied an optimized pipeline for MAGs reconstruction using as input the hybrid metagenomic assembly and both sets of short and long reads. This way, we obtained a total of 66 MAGs showing ≥50% completeness and ≤10% contamination upon quality assessment using CheckM (Parks et al., 2015) from the three Antarctic sites for which both Illumina and Nanopore sequencing were performed. Taxonomic assignment of these MAGs indicated nine phyla and 42 families, being Patescibacteria, Bacteroidota, Proteobacteria, and Actinobacteriota, the groups more represented, in agreement with diversity analysis showing the dominancy of these groups in the studied soils (Figure 5). Several of these MAGs corresponded to uncultured microbial taxa, contributing to unveiling the genomic features of this less-known part of the microbial diversity from these environments. For each MAG, we performed ARG identification and calculated the ARG load, as the number of ARGs divided by the total number of predicted CDSs. We found an overall higher ARG load in members of Patescibacteria, Proteobacteria, and Actinobacteriota. Patescibacteria showed mostly genes related to antibiotic target alteration (mainly elfamycin-resistant EF-Tu and fluoroquinolone-resistant *gyrA* and gyrB) and protection (tetracycline-resistant ribosomal protection proteins). Instead, proteobacteria showed a higher proportion of genes related to antibiotic efflux and antibiotic inactivation. From this last group, we found a class-A LRA beta-lactamase and an AIM beta-lactamase from *Burkholderiaceae* and *Nevskiaceae*, respectively, a tetracycline inactivation enzyme and a streptogramin vat acetyltransferase from *Sphingomonadaceae*, and a chloramphenicol acetyltransferase (CAT) from uncultured Rickettsiales. Also, we found mechanisms for reduced permeability to antibiotics, mainly general porins with reduced permeability to beta-lactams. Among Actinobacteriota, besides a variety of efflux mechanisms, we found rifampin monooxygenase enzymes. MAGs from Bacteroidota, although showing a reduced ARG load compared to other groups, stood out for hosting a high proportion of antibiotic inactivation enzymes and genes related to reduced permeability to antibiotics, including a VEB beta-lactamase, a CAM beta-lactamase, and a subclass B3 LRA beta-lactamase, a rifampin ADP-ribosyltransferase (Arr), and a streptogramin vat acetyltransferase. All these proteins showed 50%+ identity with the matching partner from CARD database.

**Figure 5.**
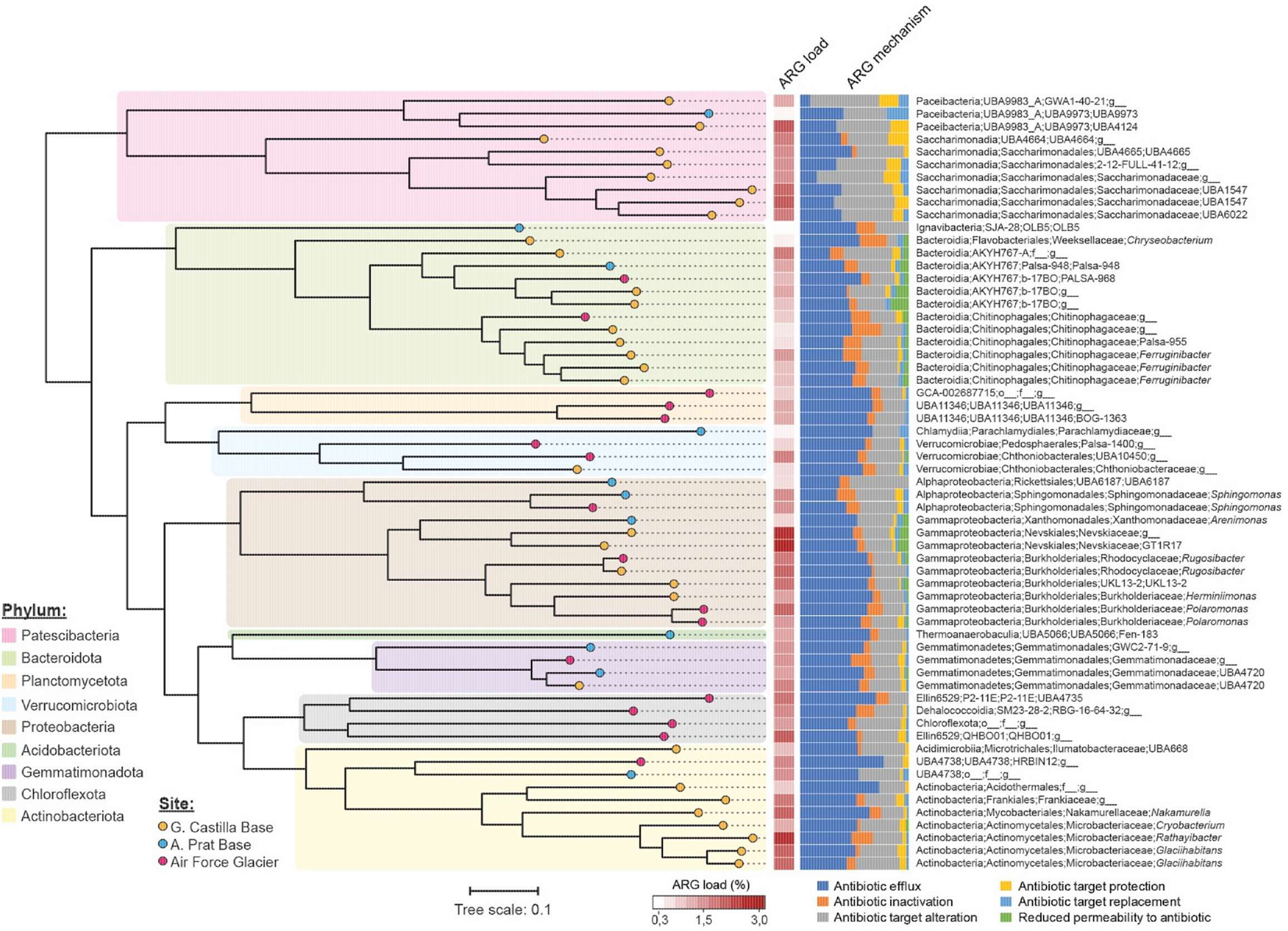
Phylogenomic diversity and ARG load of MAGs recovered from North Antarctica soil metagenomes. Sixty-one MAGs were reconstructed from the metagenomes of Air Force Glacier, G. Castilla Base, and A. Prat Base. ARG load was calculated for each MAG as the number of ARG divided by the total CDS.

### A proportion of the Antarctic ARGs are encoded inside mobile genetic elements

To evaluate if some of the ARGs identified are located inside mobile genetic elements and thus could be disseminated through horizontal gene transfer, we used the tools PlasFlow (Krawczyk et al., 2018) and Viralverify (Antipov et al., 2020), to identify putative plasmids among the assembled metagenomic contigs predicted. From 24,742 contigs harboring ARGs, 915 were predicted as plasmids by Viralverify, 3238 by PlasFlow, and 177 by both tools (Figure 6A). To minimize false positives, we focused on the last group. As a study case, we found a 15-kbp contig from *Polaromonas* harboring an OXA-like beta-lactamase that shared roughly 65% identity with OXA enzymes detected in pathogenic bacteria. Moreover, upon comparing this putative plasmid-borne Antarctic beta-lactamase with the NCBI database using BLAST, we noticed that it shares 98% identity with a class-D beta-lactamase from *Polaromonas* sp. (Accession QJS06455.1). This sequence was found to be part of a circular replicon annotated as the plasmid pA29BJ2 from *Polaromonas* sp. ANT_J29B, a strain isolated from a soil sample from Jardine Peak, King George Island (Antarctica). The metagenomic OXA beta-lactamase found harbored the conserved beta-lactamase domain (COG2602) and the amino acid residues described to be required for its characteristic activity (Paetzel et al., 2000) (Figure 6B). To have additional evidence supporting the predicted function of this Antarctic enzyme, we performed homology-based modeling of its structure using as reference three OXA beta-lactamases of known atomic-resolution structure from carbapenem-resistant *K. pneumoniae, S. enterica*, and *P. aeruginosa* (Figure 6C). From the evaluation of fifty generated models, we selected the top scored. The overall 3D structure of both the three references and the Antarctic OXA beta-lactamase were highly similar, showing 90% or more identity in relevant segments determining the characteristic alpha helices and the central beta-sheet. Moreover, the model supported a conserved disposition of the amino acid residues present in the active site, namely, Ser74, Thr75, Phe76, Lys77, Ser122, Thr123, Val124, Lys211, and Thr212. Taken together, all this evidence supports that a proportion of the North Antarctica soil resistome is located inside plasmids and potentially other kinds of mobile genetic elements. Therefore, this resistome could be transferred to pathogenic bacteria and thus act as a source of novel clinically relevant antibiotic resistance genes.

**Figure 6.**
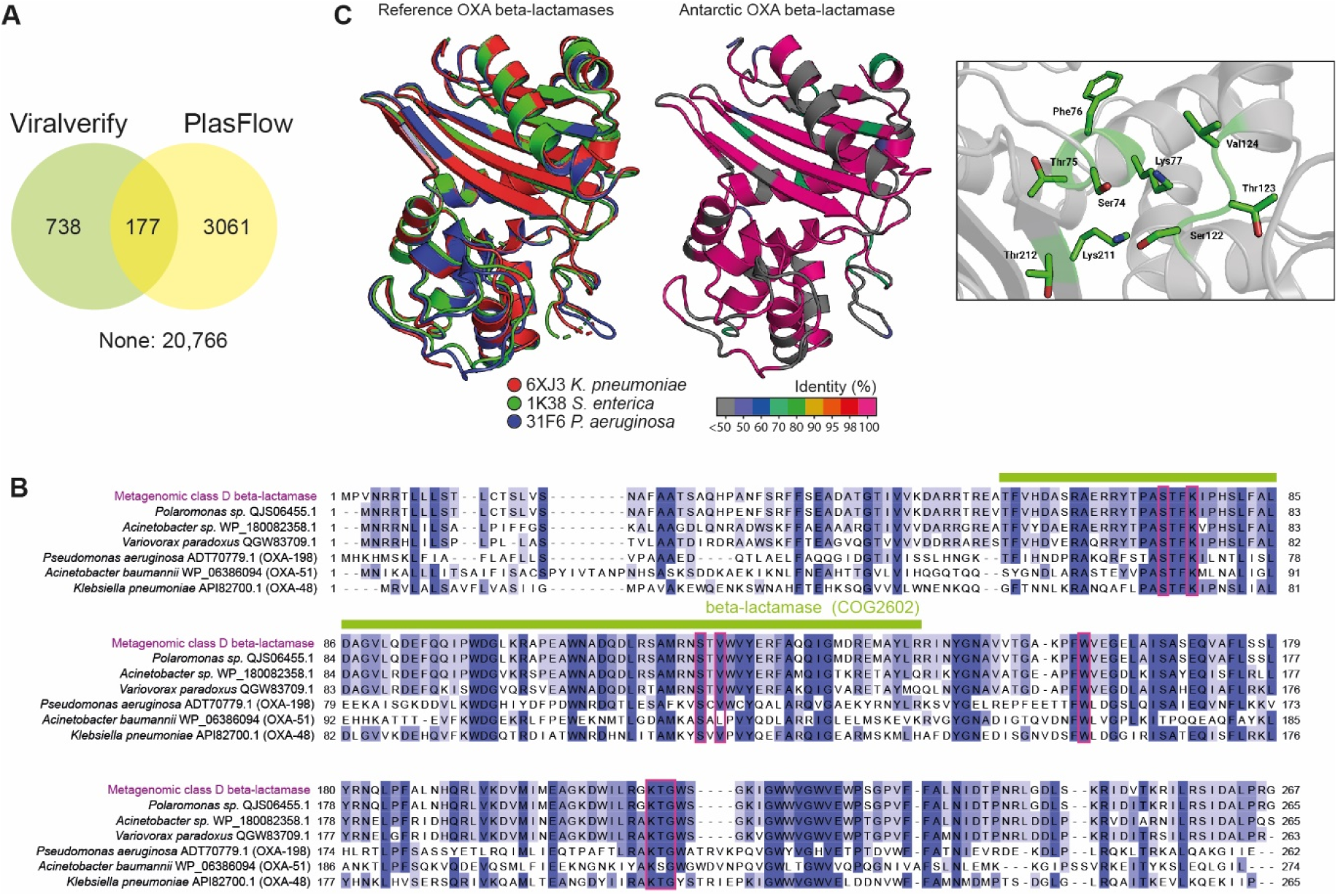
Antarctic antibiotic resistance genes found inside metagenomic contigs predicted as plasmids. (A) A total of 24,742 assembled contigs from Antarctic soil metagenomes predicted to harbor one or more antibiotic resistance genes were analysed with PlasFlow and Viralverify tools to identify contigs corresponding to plasmids. (B) Protein sequence alignment of a class-D OXA-like beta-lactamase identified in a 15 kbp contig predicted as a plasmid by both tools, and other related beta-lactamases, including three clinically relevant OXA carbapenemases from pathogenic isolates of *Pseudomonas aeruginosa, Acinetobacter bauma nii*, and *Klebsiella pneumoniae*. The green bar delimitates the detected beta-lactamase conserved domain, while red boxes indicate relevant conserved residues for the catalytic activity of this kind of enzymes. (C) Homology-based structure modeling of the Antarctic OXA-like beta-lactamase. The cartoon in the left side corresponds to the superposed structure of the three OXA enzymes used as reference, while the cartoon in the right side is the modeled Antarc tic protein, colored by the percentage identity of each region compared to the reference consensus. The inset shows a zoom of the region corresponding to the active site, indicating the most relevant amino acids described for this kind of enzymes.

## Notes

### Competing Interest Statement

The authors have declared no competing interest.

### Summary of Updates

The text was improved and shortened. We reorganized some Figures moving part of the information to the Supplementary Material. We added a section describing the reconstruction of 61 MAGs from the studied soils and the analysis of their phylogenomic relationship and resistome. Also, we added a section describing the homology-based modeling of the plasmid-borne Antarctic OXA beta lactamase from Polaromonas spp. found in our soil metagenomes, further supporting a conserved structure with OXA enzymes from multi-resistant pathogens, including the disposition of the key residues in the active site

