## Supplementary Figures and Tables for "Mapping the microbial diversity and natural resistome of North Antarctica soils"

**Running title:** Antarctic Peninsula soils natural resistome

**Keywords:** Antimicrobial resistance, natural resistome, Antarctic Peninsula, mobile genetic elements, soil microbial communities, anthropogenic intervention

**Supplementary Table 1**. Location and main features of the Antarctic sites sampled in this study.

| **Macrozon**e | Site | GPS coordinates | Relative humidity | Soil Tº | Ambient Tº | Height above sea level |
|---|---|---|---|---|---|---|
| King George Island | Ecology Glacier (NI) | S 62º 10' 16.5'' W 58º 28' 17.5'' | 63% | 1,8ºC | NR | 3 MSL |
| King George Island | Henryk Arctowski Base (H) | S 62º 9' 34.4'' W 58º 28' 18.9'' | 64% | 3,9ºC | NR | 2 MSL |
| Greenwich Island | Air Force Glacier (NI) | S 62º 28' 50'' W 59º 38' 41.5'' | 59% | 3ºC | 3,5ºC | 81 MSL |
| Greenwich Island | Arturo Prat Base (H) | S 62º 28' 44.1'' W 59º 39' 41.8'' | 74% | 2,8ºC | 2,8ºC | 0,5 MSL |
| Barrientos Island | Central area (NI) | S 62º 24' 28.2'' W 59º 44' 59.6'' | 60% | 2,4ºC | 3,3ºC | 10 MSL |
| Doumer Island | South Bay (NI) | S 64º 52' 18.6'' W 63º 33' 28.1'' | 58% | 1ºC | 1,6ºC | 80 MSL |
| Doumer Island | Yelcho Base (H) | S 64º 52' 33.3'' W 63º 35' 3.6'' | 64% | 1,5 ºC | 3,5ºC | 11 MSL |
| Deception Island | Hills behind Fumarola Bay (NI) | S 62º 57' 57,8'' W 60º 42' 51.9'' | 60% | 5,7 ºC | 4,1ºC | 20 MSL |
| Deception Island | Gabriel de Castilla Base (H) | S 62º 58' 37.3'' W 60ºC 40' 29.3'' | 66% | 5,2ºC | 4,0ºC | 10 MSL |
| Robert I Island | Coppermine Peninsula (NI) | S 62º 22' 32.7'' WO 59º 42' 44.8'' | 57% | 3,6ºC | 3,1ºC | 62 MSL |
| Robert I Island | Luis Risopatron Refugee (H) | S 62º 22' 42.8'' WO 59º 42' 3.2'' | 60% | 5,2 ºC | 3,8 ºC | 18 MSL |

**Supplementary Table 2.** Number of Antarctic isolates subjected to antimicrobial sensitivity tests per sampled zone.

| Area | Type of area | Number of isolates |
|---|---|---|
| Gabriel de Castilla base | Humanized | 35 |
| Luis Risopatron refuge | Humanized | 30 |
| Yelcho base | Humanized | 28 |
| Henryk Arctowski base | Humanized | 20 |
| Coppermine Peninsula | Non-intervened | 20 |
| South Bay | Non-intervened | 5 |
| Fumarola Bay | Non-intervened | 47 |
| Ecology glacier | Non-intervened | 44 |

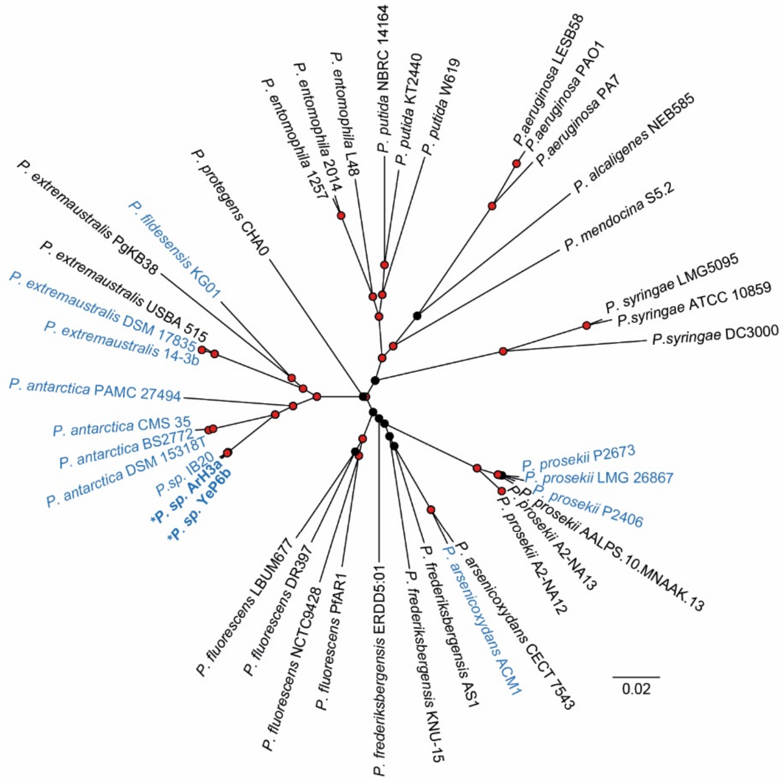


**Supplementary Figure 1. Phylogenetic relationships among the Antarctic multiresistant isolates *Pseudomonas* ArH3a and YeP6b and other species from *Pseudomonas*.** For tree construction, the assembled chromosomes of the two multiresistant isolates along with thirty eight other *Pseudomonas* strains were analyzed using Mashtree with 1,000 bootstrap iterations. Red circles indicate bootstrap higher or equal than 80%.

**Supplementary Table 3.** Strains from genus Pseudomonas included in the phylogenomic analysis of the Antarctic multi-resistant isolates *Pseudomonas* spp. YeP6b and ArH3a.

| Assembly | Reference? | Reported species | Reported strain | Antarctic? |
|---|---|---|---|---|
| GCA_000006765.1 | + | *P. aeruginosa* | PAO1 | - |
| GCA_000017205.1 | - | *P. aeruginosa* | PA7 | - |
| GCA_000026645.1 | - | *P. aeruginosa* | LESB58 | - |
| GCA_000412675.1 | + | *P. putida* | NBRC 14164 | - |
| GCA_000007565.2 | - | *P. putida* | KT2440 | - |
| GCA_000019445.1 | - | *P. putida* | W619 | - |
| GCA_001647715.1 | + | *P. antarctica* | PAMC 27494 | + |
| GCA_010634845.1 | - | *P. antarctica* | CMS 35 | + |
| GCA_900103795.1 | - | *P. antarctica* | BS2772 | + |
| GCA_900624995.1 | - | *P. antarctica* | DSM 15318T | + |
| GCA_000026105.1 | + | *P. entomophila* | L48 | - |
| GCA_003940785.1 | - | *P. entomophila* | 2014 | - |
| GCA_003940825.1 | - | *P. entomophila* | 1257 | - |
| GCA_000242115.2 | - | *P. extremaustralis* | 14-3 substr. 14-3b | + |
| GCA_001050345.1 | - | *P. fildesensis* | KG01 | + |
| GCA_900102035.1 | + | *P. extremaustralis* | DSM 17835 | + |
| GCA_900167635.1 | - | *P. extremaustralis* | USBA 515 | - |
| GCA_008692105.1 | - | *P. extremaustralis* | PgKB38 | - |
| GCA_004135995.1 | - | *P. arsenicoxydans* | ACM1 | + |
| GCA_900103875.1 | + | *P. arsenicoxydans* | CECT 7543 | - |
| GCA_900636825.1 | - | *P. fluorescens* | NCTC9428 | - |
| GCA_902825215.1 | - | *P. fluorescens* | PfAR1 | - |
| GCA_004683905.1 | - | *P. fluorescens* | LBUM677 | - |
| GCA_010448615.1 | + | *P. fluorescens* | DR397 | - |
| GCA_001874645.1 | - | *P. frederiksbergensis* | ERDD5:01 | - |
| GCA_001952935.1 | - | *P. frederiksbergensis* | AS1 | - |
| GCA_002355315.1 | - | *P. frederiksbergensis* | KNU-15 | - |
| GCA_000007805.1 | - | *P. syringae* | DC3000 | - |
| GCA_001482725.1 | - | *P. syringae* | ATCC 10859 | - |
| GCA_003047185.1 | - | *P. syringae* | LMG5095 | - |
| YeP6b (this study) | - | *P. spp.* | YeP6b | + |
| ArH3a (this study) | - | *P. spp.* | ArH3a | + |
| GCF_002263605.1 | - | *P. antarctica* | IB20 | + |
| GCA_003122265.1 | - | *P. prosekii* | P2406 | + |
| GCA_003122305.1 | - | *P. prosekii* | P2673 | + |
| GCA_003671865.1 | - | *P. prosekii* | A2-NA12 | - |
| GCA_003671895.1 | - | *P. prosekii* | A2-NA13 | - |
| GCA_008271605.1 | - | *P. prosekii* | AALPS.10.MNAAK.13 | - |
| GCA_900105155.1 | + | *P. prosekii* | LMG 26867 | + |
| GCA_001597285.1 | + | *P. alcaligenes* | NEB 585 | - |
| GCA_000397205.1 | + | *P. protegens* | CHA0 | - |
| GCA_000733715.2 | + | *P. mendocina* | S5.2 | - |

**Supplementary Table 4.** Genomic features of the Antarctic multi-resistant isolates *Pseudomonas* spp. YeP6b and ArH3a.

| Isolate | Total genome size | %GC | Plasmids | Total CDS | tRNAs | rRNAs | Prophages |
|---|---|---|---|---|---|---|---|
| *P. spp.* ArH3a | 6,774,179 | 60 | 1 (2.6 kbp) | 6,338 | 71 | 19 | 8 |
| *P. spp.* YeP6b | 6,664,416 | 60 | - | 6,141 | 67 | 19 | 7 |

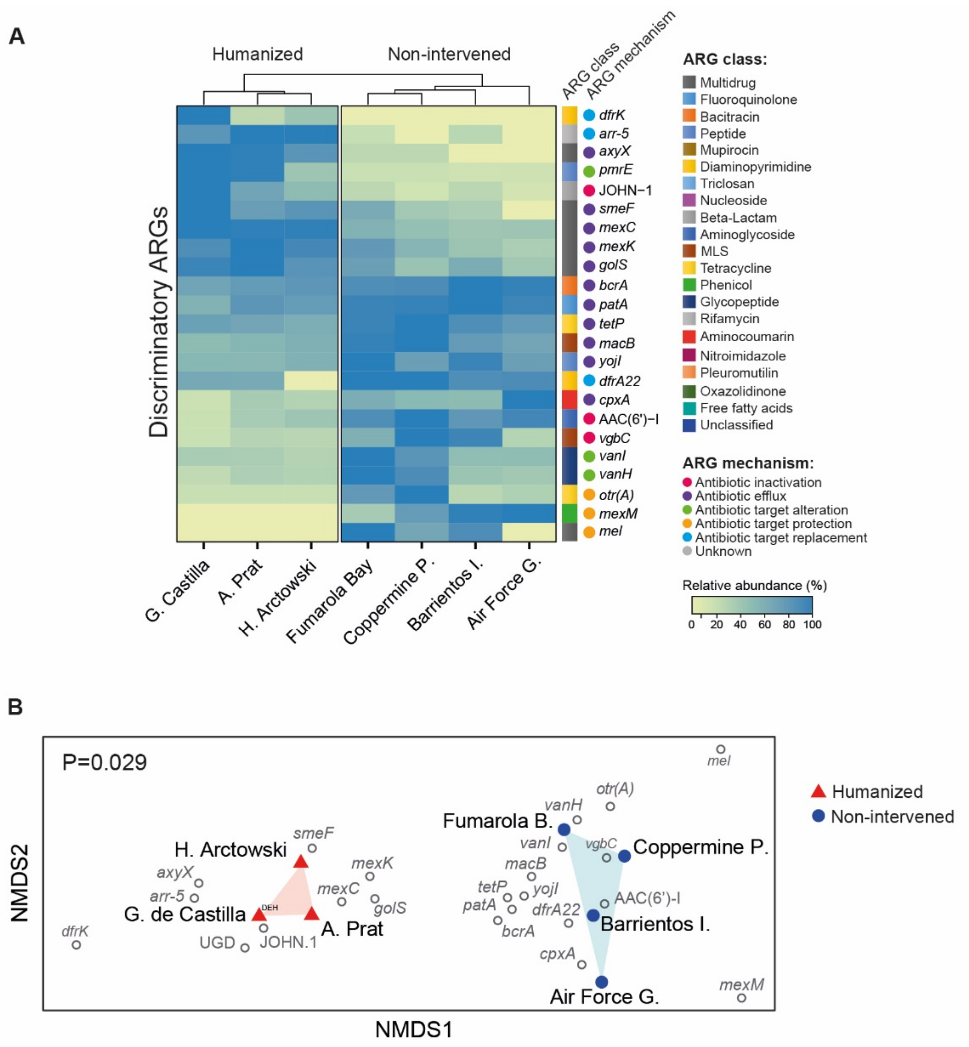


**Supplementary Figure 2. ARGs with contrasting abundance among humanized and non-intervened zones.** (A) Heatmap showing the relative abundance of discriminant ARGs identified with the extrARG tool. (B) Non-metric multidimensional scaling analysis based on the abundance dissimilarity of the discriminatory ARGs identified among the different sites (shown in gray). The P-value was calculated from a PERMANOVA analysis, supporting separate clustering of humanized and non-intervened sites.

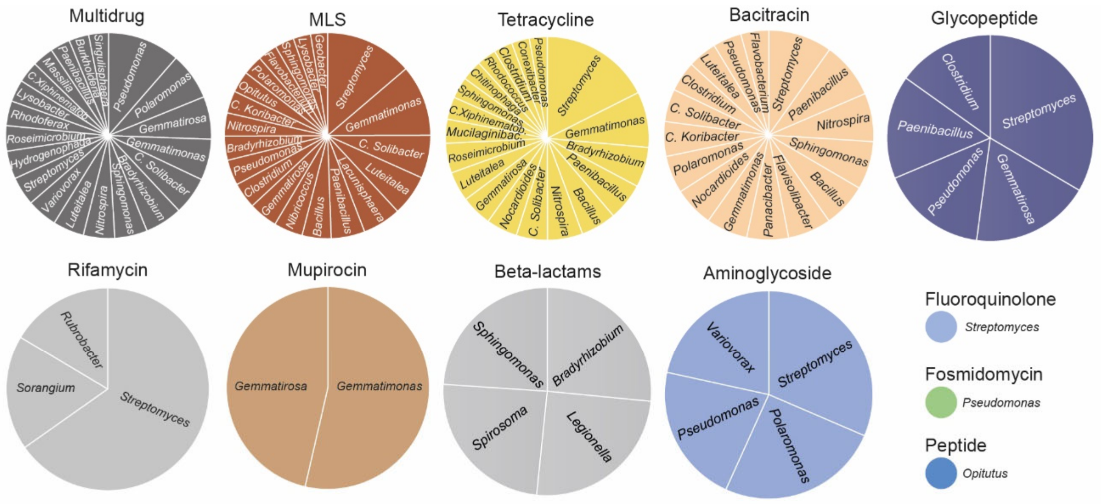


**Supplementary Figure 3. Taxonomic assignment of the Antarctic soil metagenomic reads mapping to ARGs.** Kaiju taxonomic asignment of the reads mapped to the sum of ARGs detected for each drug class.

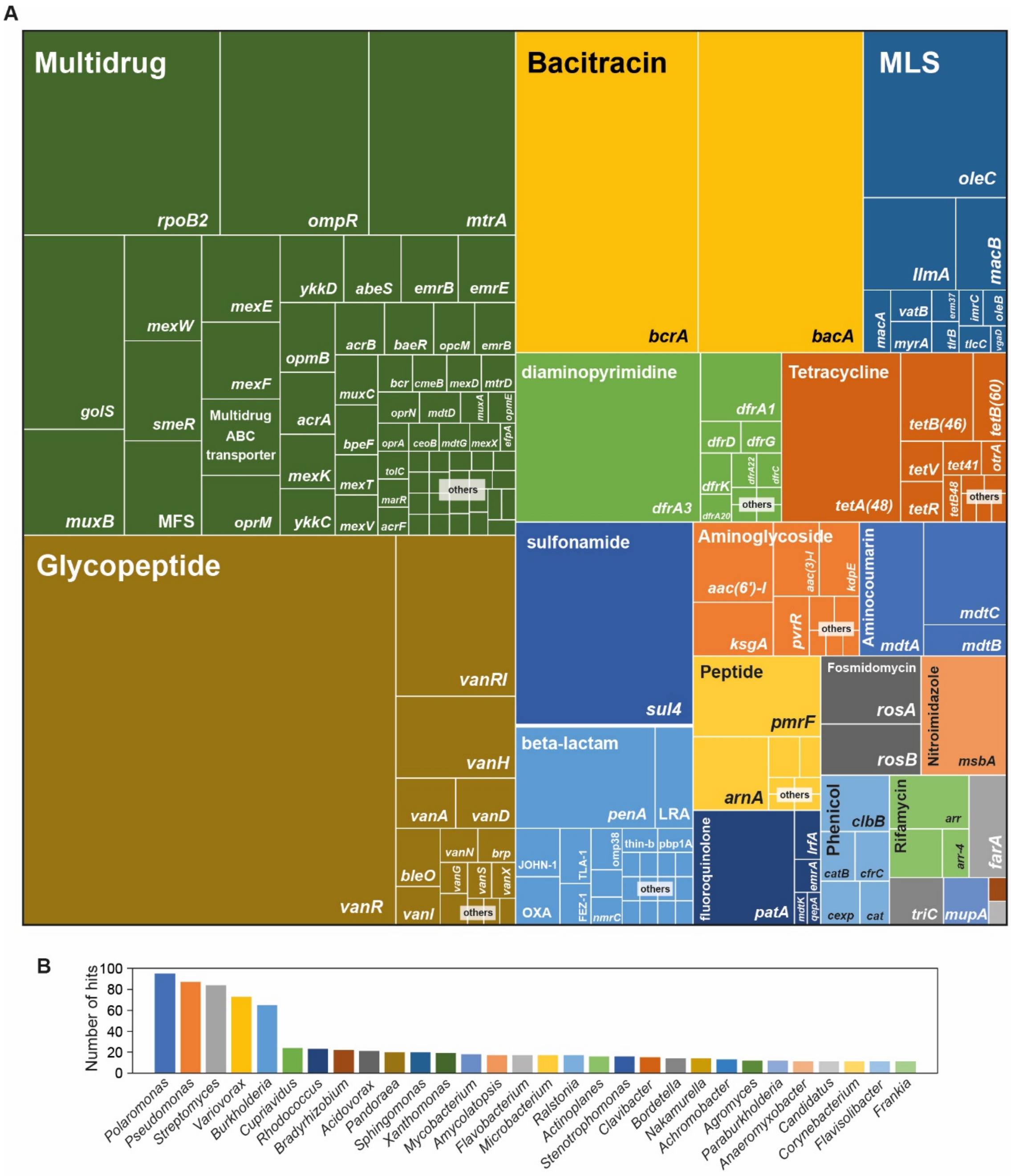


**Supplementary Figure 4. Antibiotic resistance gene families found in assembled Antarctic soil metagenomes.** (A) Treemap showing the proportion of hits found for each resistance gene family and drug class. (B) Number of contigs carrying one or more ARG that were assigned to the listed bacterial genera.
